# Distinctive Lacritin Cleavage-Potentiated Bactericidal Alteration of the *P. aeruginosa* Transcriptome

**DOI:** 10.64898/2026.03.17.712309

**Authors:** Mohammad Sharifian Gh, Fatemeh Norouzi, Gordon W. Laurie

**Affiliations:** Departments of Cell Biology, University of Virginia, Charlottesville VA, USA; Biomedical Engineering, University of Virginia, Charlottesville VA, USA; Ophthalmology, University of Virginia, Charlottesville VA, USA

**Keywords:** lacritin, N-104, bactericidal, *P. aeruginosa*, transcriptome, RNAseq, cornea

## Abstract

Lacritin is a tear, saliva, plasma and cerebrospinal fluid glycoprotein with broad polypharmacology. Its protease generated N-104 proteoform endogenous in tears (and likely elsewhere) is bactericidal and synergizes with the tear thrombin peptide GKY20. In the pathogenic and multidrug resistant PA14 strain of *P. aeruginosa*, we recently discovered that N-104 at subinhibitory and inhibitory concentrations binds to the outer-membrane lipoprotein YaiW to gain access to the periplasm where it targets and inhibits the inner-membrane ferrous iron transporter FeoB (of FeoABC) as well as PotH, a subunit of the polyamine transporter PotFGHI. Further, PA14 gene expression shifts toward anaerobic respiratory pathways. Here we report N-104-associated transcriptional changes pointing to a reduction in: (i) virulence, (ii) fitness, (iii) metabolism, (iv) stress response, (v) proteostasis, (vii) quorum sensing, and (viii) survival under anaerobic conditions. Upregulated genes are directed towards enhancing PA14: (i) multidrug and (ii) tellurite efflux, coupled with a seemingly PA14 survival attempt at (iii) anaerobic respiration, (iv) translational fidelity and (v) metabolism. The overlap with aminoglycosides (4.3%), β-lactams (0%), cyclic peptides (2.5%), fluoroquinilones (0%) and macrolide (1.9%) classes of antibiotics in *P. aeruginosa* was minimal. Thus, N-104 appears to widely perturb PA14 fundamental processes in a distinctive manner.

## 1.0 Introduction

Lacritin was originally discovered out of an unbiased biochemical screen to address the biological basis of dry eye ocular surface diseases (1). Of its several α-helices (Fig. 1A), the largely non-glycosylated (2) C-terminal amphipathic α-helix is responsible for all known lacritin activities (3–9) - including a bactericidal activity that is constitutively processed from lacritin into basal tears (9, 10). The active fragment is ‘N-104’ which comprises lacritin’s fifteen C-terminal amino acids (11, 12) (Fig. 1A). N-104 resistance screening of the 3884 Keio *E. coli* K-12 collection together with N-104 affinity screening of bacterial lysate facilitated the partial elucidation of its non-lytic killing pathway (12) in the multidrug resistant and pathogenic PA14 strain of *P. aeruginosa*. N-104 associates with outer membrane lipoprotein YaiW to enter the periplasm where it targets and inhibits the FeoB ferrous iron transporter (of FeoABC) and PotH of the PotFGHI polyamine transporter. Moreover, N-104 binds and robustly synergizes with the tear thrombin fragment GKY20 (outer membrane disruption) to kill multiple multidrug resistant clinical isolates under physiological conditions (12). N-104 is unique in its capacity to inhibit more than one transporter.

**Fig. 1.**
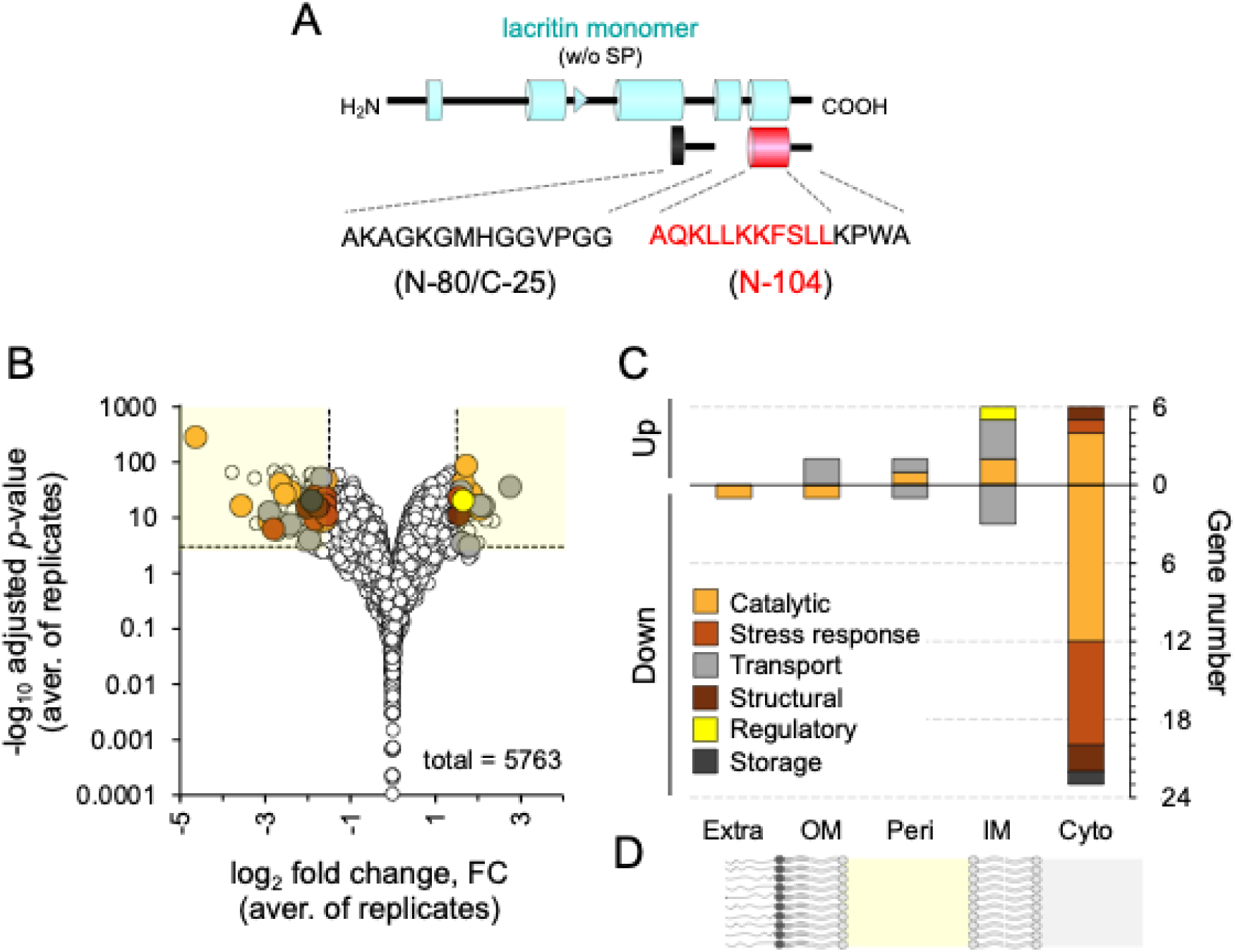
(A) Schematic representation of the secreted lacritin monomer, with blue rectangles denoting either experimentally confirmed (C-terminal half) or PSIPRED-predicted α-helices, as well as a predicted β-strand (arrowhead). Proteolytic processing yields the bactericidal proteoform N-104 (‘lacking N-terminal 104 residues’). N-80/C-25 synthetic peptide lacks the N-104 bactericidal domain and thus serves as a negative control (12). (B) Volcano plot of genes down- (left) or up- (right) regulated by N-104 relative to N-80/C-25. Circle coloring is as per C. Figure built on (12) Fig. 8A, but with different data highlighted. (C) Gene categories down- and upregulated organized by cellular location (Extra, extracellular; OM, outer membrane; Peri, periplasm; IM, inner membrane; and Cyto, cytoplasm, as schematically represented in (D).

**Fig. 2.**
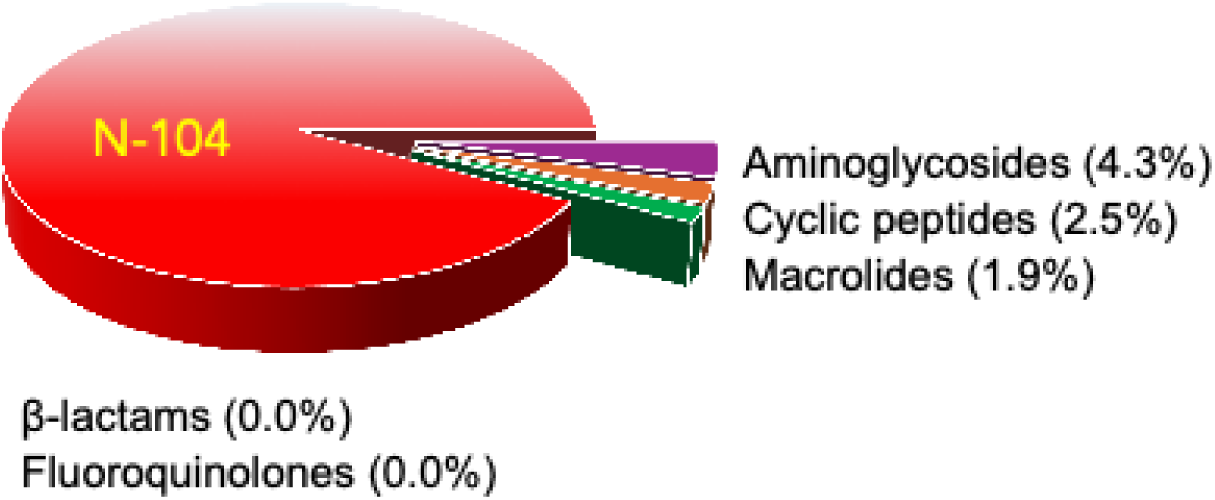
Pie chart of the relative number of N-104 altered *P. aeruginosa* genes also altered in *P. aeruginosa* by each of five classes of antibiotics.

This mechanistic approach included RNAseq that at a high level revealed a pattern of N- 104 down- and upregulation of several PA14 genes necessary for aerobic and anaerobic respiration, respectively (12). Working initially from the same data set, we now explore, elucidate and seek validation of the numerous other N-104 triggered alterations with relevance extracellularly, on the outer membrane, periplasm, inner membrane or in the cytoplasm. In keeping with its YaiW/FeoB and PotH method of action, these alterations distinguish N-104 from FDA approved antibiotics. Our goal is to identify antibiotics with new mechanisms of action which could be used to address the problem of antimicrobial resistance, particularly in multi-drug resistant *P. aeruginosa*, *A. baumannii*, and *Enterobacteriaceae* all of which are Gram-negative, and opportunistic pathogens associated with high mortality rates. PA14 as a multidrug resistant and pathogenic strain of *P. aeruginosa* serves as an excellent model for these studies (14, 15).

## 2.0 Materials and Methods

RNAseq data from our recent study (12) (NCBI GEO DataSets: Accession ID: GSE253123) was further analyzed. Rather than our earlier overview of RNAseq gene changes associated anerobic versus aerobic respiration, here we analyzed in depth and with validation all other changes in gene expression – excluding those noted in (12). From (12), overnight cultures of PA14 of which an aliquot grown in LB broth to an OD_600_ of 0.5 was pelleted, reconstituted in 1 mM Na_2_HPO_4_, 0.18 mM KH_2_PO_4_, 0.27 mM KCl, 13.7 mM NaCl (pH 7.2) and then either left untreated or treated with 20 µM N-104 or negative control N- 80/C-25 for 45 min at 35°C. N-80/C-25 is ten amino acids N-terminal to the bioactive N-104 domain (Fig. 1A) and is of similar length and isoelectric point, thus serving as an excellent negative control. We also included a 5 µM N-104 treatment arm and untreated controls. Lysozyme and SDS facilitated lysis yielding RNA’s of very high quality and integrity that were processed by UVA’s Genome Analysis and Technology Core for RNAseq and analyzed by UVA’s Bioinformatics Core. Three biological replicates were performed for all experiments. We focused on genes with an adjusted *p*-value below 0.001 (log_10_ adjusted *p*-value > 3) and a fold change greater than 2.83 (log_2_ fold change > 1.5 or < −1.5) versus the 20 µM N-80/C-25 negative control. Assignment of subcellular locations was from UniProt. Analysis tools were as per (12).

## 3.0 Results

RNAseq offers a snapshot of PA14 cellular physiology and gene regulation, although often with the assumption that mRNA and protein levels are aligned (16, 17). Among 5763 RNAseq detected genes, expression of 162 were subject to N-104 down- (118) or up-regulation (44) relative to equimolar amounts of negative control N-80/C-25 (adjusted *p* < 0.001; fold change > 2.83; Fig. 1B, C; Fig. S1)). Here we focus on those with characterized protein products whether predicted to be extracellular (‘Extra’), in the outer membrane (‘OM’), periplasmic (‘Peri’), associated with the inner membrane (‘IM’) or cytoplasmic (‘Cyto’; Fig. 1C, D). Each is presented in order of decreasing fold change (Table 1). Functions of those with currently unknown location or alternatively are uncharacterized are documented respectively in Tables 2 and 3.

**Table 1:** *P. aeruginosa* PA14 fold change in gene expression after treatment with N-104 relative to negative control N-80/C-25 showing protein products of known location and functions.

| PA14<br>gene locus | PAO1<br>homolog | FC | Protein<br>product | Location | PA14<br>gene locus | PAO1<br>homolog | FC | Protein<br>product | Location |
| --- | --- | --- | --- | --- | --- | --- | --- | --- | --- |
| <u>Downregulated</u> |  |  |  |  | <u>Upregulated</u> |  |  |  |  |
| PA14_48060 | PA1249 | -3.0 | AprA | Extra | PA14_24360 | - | 6.8 | - | OM |
| PA14_40290 | PA1871 | -5.3 | LasA | OM | PA14_60820 | PA4597 | 3.1 | OprJ | OM |
| PA14_40390 | PA1863 | -3.2 | ModA | Peri | PA14_19500 | PA3449 | 4.4 | SsuA | Peri |
| PA14_63800 | PA4825 | -7.5 | MgtA | IM | PA14_40040 | PA1893 | 3.6 | QuiP | Peri |
| PA14_02330 | PA0184 | -5.3 | AtsC | IM | PA14_13800 | PA3874 | 4.5 | NarH* | IM |
| PA14_02500 | PA0198 | -5.3 | ExbB1* | IM | PA14_03680 | PA0282 | 4.2 | CysT | IM |
| PA14_01300 | PA0106 | -4.3 | CoxA* | IM | PA14_13830 | PA3872 | 3.8 | NarI* | IM |
| PA14_12940 | PA3937 | -3.9 | TauB | IM | PA14_13780 | PA3875 | 3.5 | NarG* | IM |
| PA14_01290 | PA0105 | -2.9 | CoxB* | IM | PA14_53050 | PA0866 | 3.5 | AroP2 | IM |
| PA14_19590 | - | -24.9 | SsuF | Cyto | PA14_47040 | PA1331 | 3.4 | YegH | IM |
| PA14_34750 | PA2310 | -11.8 | TauD | Cyto | **PA14_12060** | PA4003 | 3.1 | PbpA | IM |
| PA14_19560 | PA3444 | -7.5 | SsuD | Cyto | PA14_20230 | PA3391 | 3.1 | NosR | IM |
| PA14_23680 | PA3126 | -7.0 | IbpA | Cyto | PA14_72180 | - | 3.0 | TerC | IM |
| PA14_06600 | PA0506 | -6.3 | FadE1 | Cyto | PA14_13810 | PA3873 | 5.2 | NarJ* | Cyto |
| PA14_02310 | PA0183 | -5.8 | AtsA | Cyto | PA14_19530 | PA3446 | 4.1 | SsuE | Cyto |
| PA14_24170 | PA3092 | -4.4 | FadH1* | Cyto | PA14_13260 | PA3915 | 4.0 | MoaB1* | Cyto |
| PA14_24650 | PA3049 | -4.1 | Rmf | Cyto | PA14_13850 | PA3870 | 3.8 | MoaA1 | Cyto |
| PA14_42860 | PA1673 | -3.8 | Mhr | Cyto | PA14_51680 | PA0975 | 3.3 | QueE | Cyto |
| PA14_43850 | PA1596 | -3.6 | HtpG | Cyto | PA14_13280 | PA3914 | 3.3 | MoeA1* | Cyto |
| PA14_41440 | PA1789 | -3.5 | UspA | Cyto | **PA14_15990** | PA3743 | 3.3 | TrmD | Cyto |
| PA14_01030 | PA0085 | -3.4 | Hcp1 | Cyto | PA14_27370 | PA2840 | 2.9 | DeaD | Cyto |
| PA14_71630 | PA5427 | -3.4 | AdhA | Cyto | **PA14_62780** | - | 2.9 | RimP | Cyto |
| PA14_54590 | PA0750 | -3.3 | UnG | Cyto |  |  |  |  |  |
| PA14_25180 | PA3006 | -3.3 | PsrA | Cyto |  |  |  |  |  |
| PA14_32985 | - | -3.3 | GcvH2 | Cyto |  |  |  |  |  |
| PA14_66460 | PA5027 | -3.2 | UspO | Cyto |  |  |  |  |  |
| PA14_21220 | PA3309 | -3.1 | UspK | Cyto |  |  |  |  |  |
| PA14_04650 | PA0355 | -3.1 | Pfpl | Cyto |  |  |  |  |  |
| PA14_56590 | PA4352 | -3.0 | UspN | Cyto |  |  |  |  |  |
| PA14_26140 | PA2931 | -3.0 | CifR | Cyto |  |  |  |  |  |
| PA14_33000 | - | -3.0 | GcvP2 | Cyto |  |  |  |  |  |
| PA14_60190 | PA4542 | -3.0 | ClpB | Cyto |  |  |  |  |  |
| **PA14_62990** | PA4762 | -2.9 | GrpE | Cyto |  |  |  |  |  |

**Bold, non-italic:** Core Essential gene (97, 98, 116–118)**Bold, italic:** Conditionally Essential gene (97)
FC, fold change
Extra, Extracellular; OM, Outer membrane; Peri, Periplasm; IM, Inner membrane; Cyto, Cytoplasmic; Unkn, Unknown.
\*Mentioned in (12)

**Table 2:** *P. aeruginosa* PA14 fold change in gene expression after treatment with N-104 relative to negative control N-80/C-25 showing protein products of unknown location with known or putative (UniProt) functions.

| PA14<br>gene locus | PAO1<br>homolog | FC | Protein<br>product | Function | Ref. |
| --- | --- | --- | --- | --- | --- |
| <u>Downregulated</u> |  |  |  |  |  |
| PA14_02420 | PA0193 | -13.7 | AtsK | Alkylsulfatase | - |
| PA14_19910 | PA3416 | -5.3 | PdhB | 2-oxoisovalerate dehydrogenase subunit | - |
| PA14_26130 | PA2932 | -5.1 | MorB | Morphinone reductase | - |
| PA14_34270 | PA2350 | -5.1 | MetN1 | Methionine import ATP-binding protein | - |
| PA14_34280 | PA2349 | -4.7 | MetQ1 | ABC transporter | - |
| **PA14_71720** | PA5435 | -4.6 | OadA | Oxaloacetate decarboxylase alpha chain | - |
| **PA14_25080** | PA3014 | -4.3 | FadB | Fatty acid oxidation complex subunit alpha | - |
| PA14_61530 | PA4651 | -3.9 | CsuC | Pili assembly chaperone | - |
| PA14_27520 | PA2826 | -3.8 | Gpo | Glutathione peroxidase | - |
| PA14_45020 | - | -3.7 | GlxR | Oxidoreductase | - |
| **PA14_25880** | PA2951 | -3.6 | EtfA | Electron transfer flavoprotein | - |
| PA14_07650 | PA0586 | -3.4 | YcgB | Sporulation protein | - |
| PA14_68900 | PA5217 | -3.2 | FbpA | Transferrin | (119) |
| PA14_05770 | PA0441 | -3.1 | DhT | D-hydantoinase/dihydropyrimidinase | - |
| PA14_49080 | PA1187 | -3.1 | LcaD | Acyl-CoA dehydrogenase | - |
| PA14_01620 | PA0132 | -3.0 | AptA | Adenine phosphoribosyl transferase | - |
| **PA14_31290** | PA2570 | -3.0 | Pa1L | Galactophilic lectin | - |
| **PA14_25860** | - | -3.0 | EtfB | Electron transfer flavoprotein | - |
| PA14_07680 | PA0588 | -2.9 | PrkA | AAA domain-containing protein | - |
| PA14_03610 | - | -2.9 | YcaL | Zn-dependent protease with chaperone function | - |
| <u>Upregulated</u> |  |  |  |  |  |
| PA14_47100 | PA1326 | 3.5 | IlvA2 | L-threonine dehydratase | - |
| PA14_03700 | PA0283 | 3.2 | Sbp | Sulfate-binding protein | - |
| PA14_47110 | PA1325 | 3.2 | YybH | DUF4440 domain-containing protein | - |
| PA14_66140 | PA5002 | 3.1 | DnpA | PIG-L superfamily de-N-acetylase | (120) |
| PA14_22600 | - | 2.9 | GrtA | Glycosyl transferase | - |

**Bold, non-italic:** Core Essential gene (97, 98, 116–118)
**Bold, italic:** Conditionally Essential gene (97)
FC, fold change

**Table 3:**
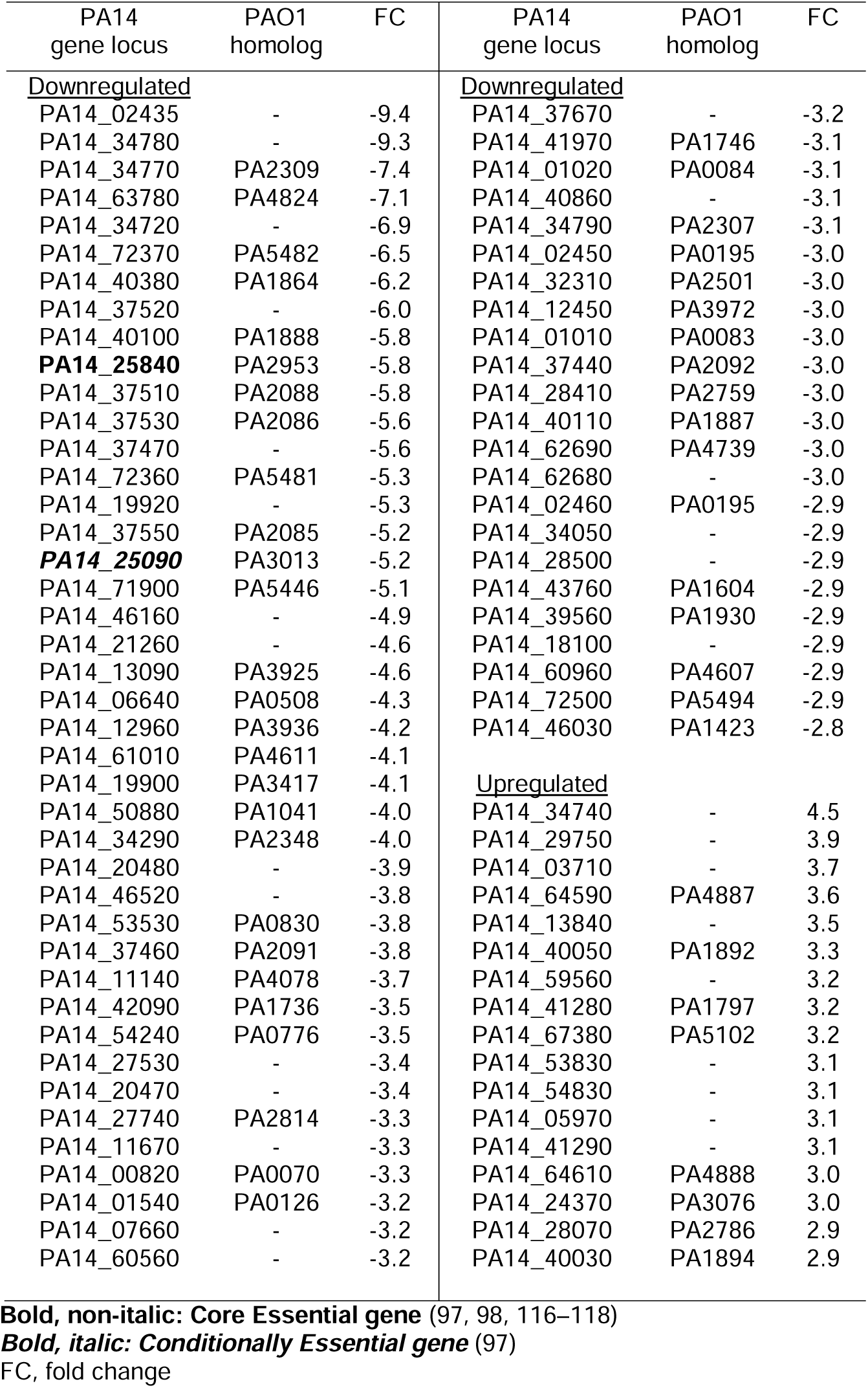
*P. aeruginosa* PA14 fold change in gene expression after treatment with N-104 relative to negative control N-80/C-25 showing uncharacterized proteins with unknown location.

| PA14<br>gene locus | PAO1<br>homolog | FC | PA14<br>gene locus | PAO1<br>homolog | FC |
| --- | --- | --- | --- | --- | --- |
| <u>Downregulated</u> |  |  | <u>Downregulated</u> |  |  |
| PA14_02435 | - | -9.4 | PA14_37670 | - | -3.2 |
| PA14_34780 | - | -9.3 | PA14_41970 | PA1746 | -3.1 |
| PA14_34770 | PA2309 | -7.4 | PA14_01020 | PA0084 | -3.1 |
| PA14_63780 | PA4824 | -7.1 | PA14_40860 | - | -3.1 |
| PA14_34720 | - | -6.9 | PA14_34790 | PA2307 | -3.1 |
| PA14_72370 | PA5482 | -6.5 | PA14_02450 | PA0195 | -3.0 |
| PA14_40380 | PA1864 | -6.2 | PA14_32310 | PA2501 | -3.0 |
| PA14_37520 | - | -6.0 | PA14_12450 | PA3972 | -3.0 |
| PA14_40100 | PA1888 | -5.8 | PA14_01010 | PA0083 | -3.0 |
| **PA14_25840** | PA2953 | -5.8 | PA14_37440 | PA2092 | -3.0 |
| PA14_37510 | PA2088 | -5.8 | PA14_28410 | PA2759 | -3.0 |
| PA14_37530 | PA2086 | -5.6 | PA14_40110 | PA1887 | -3.0 |
| PA14_37470 | - | -5.6 | PA14_62690 | PA4739 | -3.0 |
| PA14_72360 | PA5481 | -5.3 | PA14_62680 | - | -3.0 |
| PA14_19920 | - | -5.3 | PA14_02460 | PA0195 | -2.9 |
| PA14_37550 | PA2085 | -5.2 | PA14_34050 | - | -2.9 |
| **PA14_25090** | PA3013 | -5.2 | PA14_28500 | - | -2.9 |
| PA14_71900 | PA5446 | -5.1 | PA14_43760 | PA1604 | -2.9 |
| PA14_46160 | - | -4.9 | PA14_39560 | PA1930 | -2.9 |
| PA14_21260 | - | -4.6 | PA14_18100 | - | -2.9 |
| PA14_13090 | PA3925 | -4.6 | PA14_60960 | PA4607 | -2.9 |
| PA14_06640 | PA0508 | -4.3 | PA14_72500 | PA5494 | -2.9 |
| PA14_12960 | PA3936 | -4.2 | PA14_46030 | PA1423 | -2.8 |
| PA14_61010 | PA4611 | -4.1 |  |  |  |
| PA14_19900 | PA3417 | -4.1 | <u>Upregulated</u> |  |  |
| PA14_50880 | PA1041 | -4.0 | PA14_34740 | - | 4.5 |
| PA14_34290 | PA2348 | -4.0 | PA14_29750 | - | 3.9 |
| PA14_20480 | - | -3.9 | PA14_03710 | - | 3.7 |
| PA14_46520 | - | -3.8 | PA14_64590 | PA4887 | 3.6 |
| PA14_53530 | PA0830 | -3.8 | PA14_13840 | - | 3.5 |
| PA14_37460 | PA2091 | -3.8 | PA14_40050 | PA1892 | 3.3 |
| PA14_11140 | PA4078 | -3.7 | PA14_59560 | - | 3.2 |
| PA14_42090 | PA1736 | -3.5 | PA14_41280 | PA1797 | 3.2 |
| PA14_54240 | PA0776 | -3.5 | PA14_67380 | PA5102 | 3.2 |
| PA14_27530 | - | -3.4 | PA14_53830 | - | 3.1 |
| PA14_20470 | - | -3.4 | PA14_54830 | - | 3.1 |
| PA14_27740 | PA2814 | -3.3 | PA14_05970 | - | 3.1 |
| PA14_11670 | - | -3.3 | PA14_41290 | - | 3.1 |
| PA14_00820 | PA0070 | -3.3 | PA14_64610 | PA4888 | 3.0 |
| PA14_01540 | PA0126 | -3.2 | PA14_24370 | PA3076 | 3.0 |
| PA14_07660 | - | -3.2 | PA14_28070 | PA2786 | 2.9 |
| PA14_60560 | - | -3.2 | PA14_40030 | PA1894 | 2.9 |

**Bold, non-italic:** Core Essential gene (97, 98, 116–118)**Bold, italic:** Conditionally Essential gene (97)
FC, fold change

### 3.1 Extracellular

One extracellular protein is represented: serralysin AprA. Its mRNA is downregulated by N-104.

#### 3.1.2 Extracellular - Downregulated

i. **AprA** is a secreted alkaline metalloproteinase and virulence factor. By degrading *P. aeruginosa* flagellin that otherwise would be detected by host Toll-like receptor 5, AprA plays a key role in *P. aeruginosa* evasion of host innate immunity (18) including *P. aeruginosa* killing and subjection to neutrophil phagocytosis. It also inhibits host complement by C2 proteolysis. Less AprA as a consequence of N-104 treatment would significantly reduce PA14 virulence.

### 3.2 Outer Membrane

RNA’s coding for outer membrane protein (i) LasA is N-104 downregulated whereas outer membrane (ii) OprJ and (iii) PA14_24360 are upregulated.

#### 3.2.2 Outer Membrane – Downregulated

(i) **LasA** is an M23 metalloproteinase linked to *P. aeruginosa* virulence processes among which is the ectodomain shedding of host cell surface heparan sulfate proteoglycan syndecan-1 (19, 20). Lacritin targeting of syndecan-1 (4, 7) contributes to the regulation of basal tearing (1, 5), ocular surface autophagy in the context of inflammatory cytokines (8) and ocular surface epithelial and neuronal regeneration (21).

#### 3.2.3 Outer Membrane - Upregulated

(i) **OprJ** is a component of the MexCD-OprJ multidrug efflux system. Experimental studies in mice revealed that overexpression of MexCD-OprJ increased *P. aeruginosa* C3 opsonization thereby decreasing virulence (22). (iii) **PA14_24360** is a autotransporter domain containing protein. Autotransporters form the virulent type V secretion systems that deliver a diverse array of molecules into the extracellular region including those required for biofilm formation and adhesion (23).

#### 3.2.4 Outer Membrane - Summary

Taken together, less outer membrane LasA as a consequence of N-104 treatment would significantly reduce PA14 virulence. More OprJ may represent an attempt by PA14 to remove N-104 from the periplasm. The same might be true for PA14_24360, or alternatively it represents a compensatory survival mechanism.

### 3.3 Periplasm

The periplasm is the region between the outer and inner membranes (Fig. 1D). RNA for periplasmic: (i) ModA, a tungstate/molybdate/chromate-binding protein is downregulated, whereas those for (ii) SsuA, an aliphatic sulfonate binding protein and (iii) QuiP, the Acyl-homoserine lactone acylase are upregulated.

#### 3.3.2 Periplasm - Downregulated

(i) **ModA** is a periplasmic molybdate binding protein that assembles with ModB (molybdate transport) and ModC (ATPase). Molybdate is necessary for *P. aeruginosa* anaerobic respiration. Under conditions of low or no oxygen, periplasmic ModA is transported through the *P. aeruginosa* type VI secretion system into the extracellular region to capture molybdate. ModA-molybdate then associates with the outer membrane metalloproteinase IcmP from which molybdate enters the periplasm via ModABC (24), powered by ModC (25). A *P. aeruginosa* ModA transposon mutant was incapable of respiration and growth under conditions of low or little oxygen in the presence of nitrate as the terminal electron acceptor (26). Also, deficient were biofilm formation and virulence in rat lung infection and amoeba predation assays - even in the presence of molybdate (26).

#### 3.3.3 Periplasm - Upregulated

(ii) **SsuA** binds sulfonate, an organic form of sulfur. Sulfur as inorganic sulfate is preferred and required for *P. aeruginosa* survival and growth. Sulfate deficiency induces the expression of SsuA, SsuB and SsuC as an ABC sulfonate transporter together with SsuE and SsuD (27). The transporter acquires (SsuA), transports (inner membrane SsuB, cytoplasmic SsuC) and extracts sulfur from sulfonates via SsuD, a cytoplasmic flavin mononucleotide-dependent monooxygenase and SsuE, a cytoplasmic flavin mononucleotide reductase (28). When environmental sulfate is insufficient, gene expression of multiple different virulence factors is suppressed, including outer membrane LasA (above) (29). (iii) **QuiP** is an acyl-homoserine lactone acylase, a constituent of the ‘Ntn hydrolase family’ that hydrolyse *N*-acyl-homoserine lactones. *N*-Acyl-homoserine lactones are bacterial released quorum sensing signaling molecules used to guage bacterial population density (30). QuiP’s role is in part to reduce the level of *N*-3-oxododecanoyl-homoserine lactone signals. In the absence of QuiP and when the only carbon source is *N*-3-oxododecanoyl-homoserine lactone, *P. aeruginosa* displays defective growth (31).

#### 3.3.4 Periplasm - Summary

Taken together, less periplasmic ModA necessary for anaerobic respiration may couple with N-104 inhibition of anaerobic ferrous iron uptake (12) to suppress survival. More SsuA and QuiP may represent a PA14 survival response. For SsuA it might be a response to less inner membrane AtsC (sulfate ester transporter) and TauB (involved in taurine import – another sulfur source) – as both noted below.

### 3.4 Inner Membrane

Numerous RNA’s corresponding to inner membrane proteins are altered by N-104. Downregulated RNA’s include those for: (i) MgtA, an Mg^2+^ transporter of the P3B family of Mg^2+^ ATPases, (ii) AtsC, a sulfate ester ABC transporter, and (iii) TauB, an ATP binding protein taurine ABC transporter. Upregulated RNA’s for inner membrane proteins include those for: (iv) CysT, a sulfate transport system permease protein, (v) AroP2, an aromatic amino acid transport protein, (vi) YegH, involved in flavin adenine dinucleotide binding, (vii) PbpA (penicillin-binding protein 2), a core essential gene of the penicillin-binding protein family, (viii) NosR, a regulatory protein, and (ix) TerC, a membrane protein.

#### 3.4.2 Inner Membrane - Downregulated

Among proteins derived from RNA’s downregulated by N-104: (i) **MgtA**, as a transporter of magnesium into the cytoplasm is essential for *P. aeruginosa* fitness under conditions of low magnesium. This was elegantly demonstrated in *P. aeruginosa* coculture studies with the magnesium sequestering fungus *C. albicans* through transposon insertion mutant analysis, in keeping with heightened wild type *P. aeruginosa* MgtA expression under these conditions (32). Studies in *E. coli* and *S. typhimurium* have demonstrated a PhoQ/PhoP virulence determining system that triggers the expression of MgtA if magnesium is low and that MgtA preferentially associates with the anionic phospholipid cardiolipin in the inner membrane (33). (ii) **AtsC** is an ATP binding protein of the AtsRBC aromatic and aliphatic sulfate ester transporter in *P. aeruginosa* (34). (iii) **TauB** is the ATP binding component of the taurine ABC transporter consisting of taurine binder TauA and transmembrane permease TauC. Part of the *tau* cassette is *tauD* that codes for an α-ketoglutarate-dependent taurine dioxygenase. TauABCD were discovered out of an *E. coli* screen for genes regulated by sulfate starvation. TauABCD are essential for *E. coli* growth when the sole source of sulfur is taurine (2-aminoethanesulfonic acid) (35). Taurine is an important source of sulfur, carbon and nitrogen and energy for *P. aeruginosa* (36).

#### 3.4.3 Inner Membrane - Upregulated

Upregulated inner membrane RNA’s include those for (iv) **CysT**, that has not yet been characterized in *P. aeruginosa*. In *E. coli*, CysT is essential for the transport of both thiosulfate and sulfate (37) as part of an ABC transporter involving the thiosulfate binding protein CysP (and possibly sulfate binding SBP) together with CysW and CysA (38). (v) **AroP2** mRNA is one of four mRNA’s targeted by the two-component CbrAB carbon catabolite control system in *P. aeruginosa*. The system triggers a small RNA to alleviate repression of ‘catabolite repression control’ of mRNA expression – a regulated hierarchical nutrient uptake mechanism (39). As a consequence, *P. aeruginosa’s* capacity to survive under different nutrient conditions is enhanced. (vi) **YegH** has not yet been studied in *P. aeruginosa*. Gene Ontology of the *E. coli* YegH suggests a flavin adenine dinucleotide binding protein (39). (vii) **PbpA** is a D,D-transpeptidase involved in the assembly of peptidoglycan that helps maintain *P. aeruginosa*’s rod-shaped structure (40, 41). PbpA possesses an N-terminal membrane anchor and a large periplasmic catalytic domain with susceptibility to β-lactam inhibition (42, 43). (viii) **NosR** was originally discovered in *P. stutzeri* as a transcriptional regulator of the downstream adjacent *nosZ* gene coding for an N_2_O reductase (44). N_2_O reductase catalyzes N_2_O reduction as a final step in denitrification as a facilitator of bacterial anerobic growth. With motility essential for *P. aeruginosa* pathogenesis (45), it is of interest that a PA14 NosR transposon mutant is deficient in swarming associated with defective type IV pili formation necessary for swarming (45). Biofilm formation by the PA14 NosR mutant was also substantially deficient. The *nosR* gene contains methylation motifs and is under control of the DNA methylase M.PaeTBCFII in the cystic fibrosis derived and adapted *P. aeruginosa* strain TBCF10839. M.PaeTBCFII knockout mutants display reduced virulence when introduced to *G. mellonella* larvae (46). (ix) **TerC** has not been apparently studied in *P. aeruginosa*. The Te^R^ resistant cassette inclusive of *terC* was originally cloned from the pMER610 conjugative plasmid of an *Alcaligens* species (47, 48). TerC as the inner membrane element of a multiprotein pump exports the heavy metal tellurite, thereby offering resistance as per Fleming’s 1932 observation that potassium tellurite is antibacterial (49). Deletion of TerC in *K. pneumoniae* reduced virulence using the *Galleria mellonella* model. In *K. pneumoniae* TerC also provides resistance to manganese, zinc and phage (50). Studies in *B. subtilis* reveal an additional role for TerC in manganese metalation of exoenzymes necessary for lipoteichoic acid and protein synthesis (51).

#### 3.4.4 Inner Membrane - Summary

Taken together, less inner membrane MgtA alters the capacity of PA14 to restore sufficient intracellular magnesium thereby reducing PA14 fitness. The same may be true for intracellular sulfur with less AtsC and TauB. On the other hand, more CysT may be an attempt by PA14 to counter the diminishment of AtsC and TauB. More NosR reflects N-104’s promotion of anaerobic respiration (12). More TerC lowers PA14 tellurite sensitivity, possibly as an attempted PA14 survival mechanism.

### 3.5 Cytoplasm

Other N-104 altered RNA’s code for predicted cytoplasmic proteins, of which there are many. Downregulated RNA’s include those for: (i) SsuF, a putative molybdopterin-binding protein, (ii) taurine dioxygenase TauD, (iii) flavin mononucleotide-dependent monooxygenase SsuD, (iv) heat-shock proteins IbpA, (v) FadE1, a fatty acyl-CoA dehydrogenase, (vi) AtsA, an arylsulfatase, (vii) RmF, a ribosome modulation factor, (viii) Mhr, a hemerythrin functional at very low oxygen, (ix) HtpG, a chaperone protein, (x) UspA, universal stress protein, (xi) Hcp1, involved in the construction of the HIS-I type VI secretion apparatus, (xii) AdhA, an NAD-linked alcohol dehydrogenase, (xiii) UnG, uracil-DNA glycosylase, (xiv) PsrA, a transcriptional regulator, (xv) GcvH2, a glycine cleavage system protein, (xvi, xvii) UspO and UspK, universal stress proteins, (xviii) PfpI, a methylglyoxalase, (xix) UspN, a universal stress protein, (xx) CifR, a TetR-family transcriptional repressor, (xxi) GcvP2, a glycine cleavage system protein, (xxii) ClpB, a disaggregase, and (xxiii) GrpE, a heat shock protein cofactor. Upregulated RNA’s for cytoplasmic proteins include those for: (xxiv) SsuE, a flavin mononucleotide reductase, (xxv) MoaA1, GTP 3’,8-cyclase (also known as molybdenum cofactor biosynthesis protein A1), (xxvi) QueE, a 7-carboxy-7-deazaguanine synthease, (xxvii) TrmD, a tRNA methyltransferase, (xxviii) DeaD, an ATP-dependent RNA helicase and (xxix) RimP, a ribosome maturation factor.

#### 3.5.2 Cytoplasm - Downregulated

Downregulated cytoplasmic RNA’s include those for: (i) **SsuF** exhibited the greatest N-104 dependent mRNA downregulation by N-104 (Table 1). Complementation of *P. putida*transposon mutants suggest that SsuF is necessary for growth on a variety of aromatic sulfonates with the exception of methionine (52). (ii) As an α-ketoglutarate-dependent dioxygenase, **TauD** catalyzes the release of sulfite from taurine via an oxygenolytic mechanism and as previously noted is part of the TauABCD transporter. TauABCD is an important supplier of sulfur, carbon and nitrogen and energy for *P. aeruginosa* (36). Inner membrane TauB is also N-104 suppressed, as noted above. In *E. coli*, expression is regulated by the HTH-type transcription factor Cbl (35). (iii) **SsuD** extracts sulfur from sulfonates in a manner dependent on cytoplasmic flavin mononucleotide reductase SsuE that for E. coli is optimal in vitro at an SsuE/SsuD molar ratio of 2.1 – 4.2 (53). Complementation of *P. putida* transposon mutants suggest that SsuD is necessary for growth on sulfonates or methionine (52). (iv) **IbpA** was first discovered as an inclusion body associated protein in *E. coli* stressed by recombinant protein production (54). IbpA is a heat shock protein that promotes cellular proteostasis by facilitating protein refolding, protein aggregate disassociation and the removal of misfolded proteins. Expression of IbpA in *P. aeruginosa* is by the heat-shock sigma factor RpoH (55). Degradation is contributed by the Lon ATP-dependent serine protease (56). IbpA is thought to contribute to oxidative stress tolerance in *P. aeruginosa*, notably when the OxyR regulatory pathway is compromised (57). (v) **FadE1** mediates the conversion of 3-(methylthio)propanoyl-CoA to 3-(methylthio)acryloyl-CoA, with strong specificity for long-chain acyl-CoAs as part of the fatty acid metabolism pathway. Fatty acids are a primary carbon source for *P. aeruginosa* in cystic fibrosis. *P. aeruginosa* lacking FadE1 displayed diminished growth on C14:0, C16:0 and C18:1 fatty acids as the sole carbon source, but not on glucose, C6:0, C8:0 or C10:0 fatty acids. Virulence in the *Galeria mellonella* infection model was also diminished (58). (vi) **AtsA**, together with N-104 downregulated inner membrane AtsC of the sulfate ester transport system AtsRBC (34) supports growth by increasing sulfur availability in a sulfate, cysteine, or thiocyanate inhibitable manner (59). In *P. aeruginosa*, the *ats* genes are considered to be an extension of the *cys* regulon regulated by cysB (34) - noting that mRNA of inner membrane CysT of the CysAWTP transporter is upregulated by N-104. (vii) **RMF** promotes dormancy by contributing (60) to the dimerization of 70S ribosomes into inactive but stable 100S ribosomes - a process known as ribosome hibernation (61). Dormant, stressed, cells reside deep in *P. aeruginosa* biofilms and display less sensitivity to the aminoglycoside tobramycin and fluoroquinolone ciprofloxacin (62). Upon addition of tobramycin to *P. aeruginosa* biofilms, the RMF gene is the most highly activated (63). RMF may also play a role in supporting dormant cell viability since membranes of *P. aeruginosa* lacking RMF are propidium iodide permeable (62). (xv) **Mhr** is regulated by Anr, the low oxygen sensor transcription factor. As a hemerythrin, Mhr binds oxygen via a di-iron center with micromolar affinity thereby contributing modestly to *P. aeruginosa* survival under low oxygen conditions and enhancing competitiveness against *E. coli*. It does so, in part, by exiting the cell for oxygen capture through the type VI secretion system. Mhr-oxygen is then retrieved by the low oxygen expressed outer membrane protein OprG (facilitated possibly by direct membrane interaction (64)) for entry into the periplasm where it binds inner membrane subunits of cytochrome C oxidase - also under Anr low oxygen regulation (65). Hemerythrins are found in all domains of life (66). (ix) **HtpG** is a homolog of heat shock protein 90 with virulence activity. Its absence in *P. aeruginosa* is associated with the apparently diminished capacity of *P. aeruginosa* supernatant to lyse boiled S. aureus cell pellet as an indication of LasA protease activity and involvement in other *P. aeruginosa* functions (67). Also, it triggers the release of inflammatory IL8 from macrophages differentiated from THP-1 human monocytes (68). (x) The universal stress protein **UspA** is also regulated by the Anr low oxygen sensor transcription factor and is not apparently required for survival. (xi) **Hcp1**, which assembles as a hexameric oligomer with a 40Å internal ring diameter to form the toxin injecting spear of the type VI secretion system. Hcp1 is expressed from the *P. aeruginosa* HIS-I virulence locus essential for lung infectivity under the respective positive and negative control of virulence regulators LadS and RetS. Hcp1 is particularly prevalent in pulmonary secretions of cystic fibrosis patients chronically infected with *P. aeruginosa* (69). Further regulated by Anr is (xii) **AdhA** that accordingly displays elevated expression when oxygen is limiting. Lacking AdhA, *P. aeruginosa* fails to grow under low or no oxygen on ethanol as the sole carbon source (70). (xiii) **UnG** is essential for genome maintenance by removing uracil from DNA that can arise by the deamination of cytosine (71). *P. aeruginosa* lacking UnG display a four-fold enhanced mutational frequency (72). Cytosine deamination is enhanced in the presence of nitric oxide. *P. aeruginosa* lacking Ung under conditions generating nitric oxide showed decreased growth and after phagocytosis are substantially less viable (73). (xiv) **PsrA** transcriptionally regulates the *P. aeruginosa* type III secretion system (74) and many other genes (75, 76). Lack of PsrA in *P. aeruginosa* is associated with reduced biofilm formation, swarming motility and attachment (76) together with reduced resistance to phagocyte-like PLB-985 cells (74). (xv) **GcvH2** is a key component of the glycine-inducible glycine cleavage system that regulates bacterial utilization of glycine as a carbon source for protein and metabolite production (77). An inhibitor of the mitochondrial glycine cleavage system is cysteamine. Cysteamine treated wild type *P. aeruginosa* and *P. aeruginosa* lacking GcvH2 fail to grow in M9 medium containing glycine as the sole carbon source (78). Also anaerobic stress induced are (xvi) **UspO** and (xvii) **UspK**. UspO is not apparently required for survival but regulated by the Anr low oxygen sensor transcription factor. UspK is required for *P. aeruginosa* survival during anaerobic long-term pyruvate fermentative growth (79, 80). (xviii) **PfpI** converts toxic methylglyoxal to lactic acid in a glutathione-dependent manner. As an electrophile, methylglyoxal can target protein amino and thiol groups, as well as guanosine in DNA. For proteins this can be inactivating and is damaging for DNA. Indeed, *P. aeruginosa* lacking PfpI revealed a significant 2-fold downregulation of eighteen proteins including several required for translational efficiency, glutathione-dependent glyoxalase and repression of the MexCD-OprJ efflux pump (81). The original description (82) of *Pfpi* as a antimutator gene whose protein product contributed to the alleviation of cellular stress has been disproven (81). Anaerobic stress induced (xix) **UspN** is required for *P. aeruginosa* survival during anaerobic stationary phase as observed in *P. aeruginosa* lacking it (79, 80). (xx) **CifR** binds and represses expression of the P. aeruginosa toxin Cif in an epoxide inhibitable manner (83). Cif is an epoxide inducible epoxide hydrolase that reduces expression of the epithelial cystic fibrosis transmembrane conductance regulator (84). Epoxides are important lipid signaling metabolites of arachidonic acid (85). (xxi) **GcvP2** is a key component of the glycine-inducible glycine cleavage system that regulates bacterial utilization of glycine as a carbon source for protein and metabolite production (77). (xxii) **ClpB** is a chaperone that enables recovery from heat-induced protein aggregation (86, 87) by cooperating with the chaperone DnaK to disaggregate and reactivate damaged proteins. (xxiii) **GrpE** is an ADP/ATP exchange factor that as studied with GrpE from other bacterial species transitions DnaK from a ADP-liganded substrate-binding high affinity state into an ATP-liganded low affinity state for substrate release – with kinetics temperature-modulated by GrpE (88). GrpE is is a *P. aeruginosa* core essential gene (Table 1).

#### 3.5.3 Cytoplasm - Upregulated

Those with predicted cytoplasmic proteins that are upregulated include: (xxiv) **SsuE** is necessary for growth on aromatic sulfonates but not methionine or aliphatic sulfonates, as per complementation of *P. putida* transposon mutants. Information of *P. aeruginosa* SsuE is lacking beyond the comparative genetic organization of the ssu operons among bacterial species (52). (xxv) **MoaA1** catalyzes the cyclization of GTP to (8S)-3’,8-cyclo-7,8-dihydroguanosine 5’-triphosphate (3’,8-cH2GTP) for conversion to cyclic pyranopterin monophosphate (‘cPMP’) by MoaC (cPMP synthase). Pyranopterin is a ring structure precursor to the biosynthesis of molybdenum cofactor (‘Moco’) (89). Biosynthesis of molybdenum cofactor is conserved among all kingdoms of life to make molybdenum available to molybdoenzymes important in carbon, nitrogen and sulfur metabolism and traceable to LUCA (90). MoaA1 in *P. aeruginosa* remains to be explored. (xxvi) **QueE** contributes to the queuosine hypermodification of tRNA’s for asparagine, aspartic acid, histidine and tyrosine in position 34 of the anticodon wobble (91). Queuosine is a 7-deazaguanosine nucleoside in most archaea, bacteria and eukarya (92). In *P. aeruginosa*, QueE behaves as a single-function biosynthetic enzyme, lacking a stress-induced secondary role in promoting cell size and division in *E. coli* (93). (xxvii) **TrmD** which catalyzes the methylation of guanosine-37 in *P. aeruginosa* tRNAs including those for leucine (CUN), proline (CCN) and arginine (CCG) thereby preventing frame-shifting (94). TrmD is conserved in archaea, bacteria and eukarya, as well as organelles (95) and is considered either essential (96) or conditionally essential (97, 98) or neither (98) in *P. aeruginosa*. (xxviii) **DeaD** in *P. aeruginosa* serves as a RNA-helicase to posttranscriptionally facilitate the translation of HTH-type transcriptional regulator ExsA, which in turn promotes the gene expression of the type III secretion system (99). (xxix) **RimP** is core essential in *P. aeruginosa* (97, 98). Mechanistic studies have been performed in E. coli and (100) *S. typhimurium* (101) revealing a critical role in 30S ribosomal subunit assembly to which it binds (100) by promoting proper folding of 16S rRNA. *E. coli* lacking *rimP* display deficient 30S ribosomal maturation associated with impaired growth at 30, 37 and 44°C.

#### 3.5.4 Cytoplasm - Summary

Taken together as a consequence of N-104 treatment, less Hcp1 and PsrA may respectively compromise assembly of the type VI and III secretion systems thereby reducing virulence. Less CifR may increase the availability of Cif to reduce the cystic fibrosis transmembrane conductance regulator. Less GcvH2, GcvP2, FadE1, AdhA, UspK and UspN may limit carbon source growth options under normoxia (GcvH2, GcvP2, FadE1) or hypoxia (AdhA) or no oxygen (UspK and UspN) conditions. Less TauD (together with less inner membrane TauB), less AtsA (together with less inner membrane AtsC), less SsuD and SsuF may respectively reduce the capacity to harvest sulfite from taurine, import sulfur and extract sulfur from sulfonates. Less ClpB may diminish recovery from heat shock. Less GrpE may alter the capacity of DnaK to function fully as a chaperone. Less RMF may reduce *P aeruginosa* viability when dormant. Less UnG or PfpI may compromise viability. More TrmD (frame-shift prevention), QueE (queuosine tRNA hypermodification, RimP (30S ribosomal maturation) and MoaA1 (Moco biosynthesis for molybdenum presentation to molybdoenzymes) and DeaD (promotes gene expression of the type III secretion system) might be a secondary cellular response to N-104 killing. N-104 compromises global metabolic flexibility and stress tolerance.

### 3.6 N-104 Comparison to Antibiotic Classes

N-104’s killing mechanism is transcriptionally extensive. Does it differ from aminoglycosides, β-lactams, cyclic peptides, fluoroquinilones and macrolide classes of antibiotics favored for *P. aeruginosa*? Differences might be expected with N-104 suppressing polyamine and ferrous iron uptake while seemingly downregulating aerobic respiration in favor of anaerobic respiration – despite ferrous iron being the preferred iron source under anaerobic conditions (12). In contrast, aminoglycosides bind 30S ribosome 16S ribosomal RNA to alter protein translational accuracy to disrupt membrane integrity and proteostasis (102), β-lactam binds *P. aeruginosa* penicillin-binding protein 3 required for peptidoglycan cross-linking (103), cyclic peptides are membrane lytic by initially disrupting the *P. aeruginosa* outer lipopolysaccharide layer (104), fluoroquinilones obstruct *P. aeruginosa* DNA integrity by inhibiting DNA gyrase and topoisomerase IV (105) and macrolides alter *P. aeruginosa* protein translation by binding the ribosomal polypeptide exit tunnel (106). We compared published *P. aeruginosa* transcriptional changes triggered by each (107–111) to that of N-104. The overlap with aminoglycosides (4.3%), β-lactams (0%), cyclic peptides (2.5%), fluoroquinilones (0%) and macrolide (1.9%) classes of antibiotics in *P. aeruginosa* was minimal. Table 4 documents similarities.

**Table 4:** *P. aeruginosa* PA14 N-104 dependent gene changes shared with aminoglycosides, β-lactams, cyclic peptides, fluoroquinilones or macrolides.

| Antibiotic Class | Antibiotic | Downregulated | Upregulated | Culture (Ref.) |
| --- | --- | --- | --- | --- |
| Aminoglycoside | Tobramycin | PA14_19900<br>PA14_19920<br>PdhB | DeaD<br>IlvA2<br>TrmD<br>RimP | planktonic PA14 (107) |
| β-lactam | Ceftazidime | - | - | planktonic PA14 (110) |
|  | Imipenem | - | - | planktonic PAO1 (121)<br>PAO1 biofilms (121) |
|  | Meropenem | - | - | PAO1 biofilms (122) |
| Cyclic peptide | Polymyxin B | - | PA14_24360 | planktonic PA14 (111) |
|  | Polymyxin E | - | OprJ<br>NosR<br>PA14_24360<br>PA14_41280 | planktonic PA14 (107) |
| Fluoroquinolone | Ciprofloxacin | - | - | planktonic PAO1 (123) |
|  | Levofloxacin | - | - | planktonic PAO1 (124)<br>planktonic PA14 (108) |
| Macrolide | Azithromycin | AprA<br>LasA<br>PfpI | - | planktonic PAO1 (109)<br>planktonic PA14 (110) |

## 4.0 Discussion

Identification of antibiotics with new mechanisms of action may help address the ongoing and pervasive threat of antimicrobial resistance, particularly in multi-drug resistant *P. aeruginosa*, *A. baumannii*, and *Enterobacteriaceae* - all of which are Gram-negative and opportunistic pathogens associated with high mortality rates and for *P. aeruginosa* also ocular surface pathology. Using a transcriptomic approach, we report that lacritin cleavage-potentiated tear bactericide N-104 broadly disables the virulent PA14 strain of *P. aeruginosa* in a manner disinct from aminoglycosides, β-lactams, cyclic peptides, fluoroquinilones and macrolide classes of antibiotics favored for the treatment of *P. aeruginosa*. It does so by transcriptionally reducing the capacity for virulence, fitness, metabolism, stress response, proteostasis and quorum sensing. It also appears to diminish survival under anaerobic conditions while paradoxically promoting anaerobic respiration at the expense of aerobic respiration (12). N-104 is a tear proteoform of lacritin, as revealed by top-down mass spectrometry and with validating bactericidal prediction and assessment of overlapping N-105, N-106 and N-107 (9, 10).

N-104 first binds the *P. aeruginosa* outer membrane lipoprotein YaiW to facilitate periplasmic entry. It then uniquely inhibits both polyamine and ferrous iron uptake by respectively targeting the inner membrane PotH subunit of the PotFGHI polyamine channel and inner membrane FeoB, a ferrous iron transporter. FeoB is the favored iron transporter under anaerobic conditions. Thus, N-104 promotes an environment suitable for FeoB while counterproductively serving as an effective FeoB inhibitor. Although *P. aeruginosa* displays a propensity for acquiring resistance against cationic antimicrobial peptides by 10 generations, none was developed against N-104 through 30 succesive generations. N-104 both binds and synergizes with tear antimicrobial peptide GKY20 to kill clinical isolates of *P. aeruginosa*, *S. marcescens* and *S. maltophilia* (12).

Human tears are rich in antimicrobial factors (112). Whole human stimulated tears have been shown to be bacteriostatic for *P. aeruginosa* strains PA103 and 6206 (113). Basal tear bactericidal activity for *P. aeruginosa* and *E. coli* was substantially immunodepleted using antibodies against lacritin’s C-terminal domain inclusive of N-104 (11). Metabolomic analysis of *E. coli* treated for 15 minutes with the larger lacritin C-terminal N-65 fragment revealed that cellular metabolic activity had been broadly compromised (11), in keeping with our N-104 triggered transcriptomic data. Both approaches also revealed evidence of cellular prosurvial strategies. Human tears collected by capillary tube without anesthesia then incubated with *P. aeruginosa* for 5 hours also broadly altered gene expression (114), including respectively 15 and 4 downregulated and upregulated genes shared with N-104. The tear activity was insensitive to boiling, but sensitive to proteinase K and enhanced *P. aeruginosa* killing activity of polymyxin, but not aminoglycoside, carbapenem, cephalosporin, rifamycin or tetracyline classes of antibiotics (114).

Taken together, N-104’s properties may offer promise for the alleviation of Gram-negative, multi-drug resistant bacterial pathogenicity of the eye and elsewhere – either alone or possibly in synergy with aminoglycosides, β-lactams, carbapenems, fluorquinolones and polymyxins as has been demonstrated for other cationic antimicrobial peptides (115).

## CRediT authorship contribution statement

**Mohammad Sharifian Gh.:** Conceptualization, Data curation, Formal analysis, Investigation, Methodology, Validation, Visualization, Writing - original draft, Writing - review & editing. **Fatemeh Norouzi:** Conceptualization, Data curation, Formal analysis, Investigation, Methodology, Validation Writing - review & editing. **Gordon W. Laurie:** Conceptualization, Formal analysis, Funding Acquisition, Methodology, Project administration, Resources, Supervision, Validation, Visualization, Writing - original draft, Writing - review & editing.

## Acknowledgments

We acknowledge UVA’s Katia Sol-Church (Genome Analysis and Technology Core) and Pankaj Kumar (Bioinformatics Core) and Robert A. Bloodgood for reviewing the manuscript.

## Data availability

Data will be made available on request.

